# Inter-species differences in response to hypoxia in iPSC-derived cardiomyocytes from humans and chimpanzees

**DOI:** 10.1101/382895

**Authors:** Michelle C. Ward, Yoav Gilad

**Affiliations:** Department of Medicine, University of Chicago, Chicago, IL, USA; Department of Human Genetics, University of Chicago, Chicago, IL, USA

## Abstract

Despite anatomical similarities, there appear to be differences in susceptibility to cardiovascular disease between primates. For example, humans are prone to ischemia-induced myocardial infarction unlike chimpanzees, which tend to suffer from fibrotic disease. However, it is challenging to determine the relative contributions of genetic and environmental effects to complex disease phenotypes within and between primates. The ability to differentiate cardiomyocytes from induced pluripotent stem cells (iPSCs), now allows for direct inter-species comparisons of the gene regulatory response to disease-relevant perturbations. A consequence of ischemia is oxygen deprivation. Therefore, in order to understand human-specific regulatory adaptations in the heart, and to potentially gain insight into the evolution of disease susceptibility and resistance, we developed a model of hypoxia in human and chimpanzee cardiomyocytes. We differentiated eight human and seven chimpanzee iPSC lines into cardiomyocytes under normoxic conditions, and subjected these cells to 6 hours of hypoxia, followed by 6 or 24 hours of re-oxygenation. We collected genome-wide gene expression data as well as measurements of cellular stress at each time-point. The overall cellular and transcriptional response to hypoxic stress is generally conserved across species. Supporting the functional importance of precise regulatory response to hypoxia, we found that genes that respond to hypoxic stress in both species are depleted for association with expression quantitative trait loci (eQTLs) in the heart, and cardiovascular-related genes. We also identified hundereds of inter-species regulatory differences in our study. In particular, *RASD1*, which is associated with coronary artery disease, is up-regulated specifically in humans following hypoxia.

## Introduction

Understanding human susceptibility to disease is a central goal of biomedical research. One approach to gaining insight into human adaptation, resistance and susceptibility to cellular stress and, ultimately disease, is to investigate the genetic, molecular, cellular and morphological differences between humans and our closest evolutionary relatives, the great apes.

Cardiovascular disease (CVD), for example, is responsible for about a third of both human and captive chimpanzee deaths (WHO; (Varki et al. 2009)). The anatomy of the healthy human and chimpanzee heart is similar; however CVD pathology differs. Chimpanzee disease is often associated with interstitial myocardial fibrosis, while human heart disease predominantly results from coronary artery artherosclerosis, leading to ischemic damage (Lammey et al. 2008; Varki et al. 2009).

A consequence of myocardial ischemia, the reduction of blood flow to the heart tissue, is oxygen deprivation. Oxygen sensing and response is an essential process across species. If the balance between anti-oxidants and pro-oxidants, such as reactive oxygen species (ROS), is decoupled, redox signaling is disrupted, and oxidative stress ensues. The heart is the most oxygen-demanding tissue in the body (Giordano 2005). Maintenance of oxygen homeostasis is essential for cardiac function as imbalance of ROS can result in myocardial infarction and heart failure. Indeed, 20-40 minutes of oxygen deprivation results in irreparable damage to the human heart (Bretschneider et al. 1975).

It is well appreciated that CVD is a complex disease with many contributing genetic and environmental factors. These effects are difficult to distinguish in *in vivo* studies within and between species because, in order to establish clear causal relationships and mechanism, directed perturbation is required. This is infeasible in humans and other apes due to practical and ethical considerations. More tractable model organisms such as mice are not optimal models of CVD given the differences in relative heart size and heart rate (Doevendans et al. 1998; Milani-Nejad and Janssen 2014). The advent of induced pluripotent stem cell (iPSC) technology now allows us to access disease-relevant cell types across human individuals and other primates, and to test the effects of controlled perturbation. We have recently established a panel of human and chimpanzee iPSC lines (Gallego Romero et al. 2015), and we have shown that cardiomyocytes derived from induced pluripotent stem cells (iPSC-CMs) can effectively model gene regulation in hearts from humans and chimpanzees (Pavlovic et al. 2018).

Cardiomyocytes make up 70-85% of the heart volume, 30-40% of the total cellular composition (Pinto et al. 2016; Zhou and Pu 2016), and are susceptible to ischemia following coronary artery occlusion. In order to gain insight into human gene regulatory adaptation in the heart, and the evolution of disease susceptibility and resistance, we developed a model of hypoxia and reoxygenation in human and chimpanzee iPSC-CMs. This cell culture-based system enables an indepth characterization of the inter-species response to, and recovery from, hypoxic stress. We can now determine intrinsic inter-species regulatory differences in response to a universal cellular stress, which could provide insight into the observed phenotypic differences in the manifestation of CVD between species.

Hypoxia induces a transcriptional response following stabilization of the HIF1 transcription factors under conditions of low oxygen (Samanta and Semenza 2017). We therefore determined both the global transcriptional response to hypoxia and re-oxygenation by RNA-seq, and the cellular response by measuring features of oxidative damage including lipid peroxidation and DNA damage, in both species. While an iPSC-derived cardiomyocyte-based system has been previously used to study the effects of hypoxia in a single human and a single rhesus macaque individual, here we use a panel of human and chimpanzee individuals to identify a set of conserved and species-specific response genes (Zhao et al. 2018). The identification of inter-species gene regulatory differences allowed us to develop hypotheses about molecular mechanisms that might explain phenotypic differences between species.

## Results

We differentiated cardiomyocytes from iPSCs of eight human and seven chimpanzee individuals, including replicate differentiations from a subset of the lines (Figure 1A, and Figure S1A). The 15 human and chimpanzee iPSC lines we used have been well characterized as described here and previously (Figure S2 and Table S1) (Gallego Romero et al. 2015; Burrows et al. 2016; Pavlovic et al. 2018; Ward et al. 2018). To increase the purity of the cardiomyocyte cultures, we used a metabolic purification step (Tohyama et al. 2013). To obtain more mature cardiomyocytes, we cultured the cells for 30 days post induction of differentiation (Chan et al. 2013; Robertson et al. 2013), subjected the cells to electrical stimulation to increase cellular elongation and improve calcium handling (Chan et al. 2013), and cultured the cells in the presence of galactose instead of glucose to shift the cells’ metabolism from fetal-associated glycolysis to adult-associated mitochondrial respiration (Rana et al. 2012).

**Figure 1:**
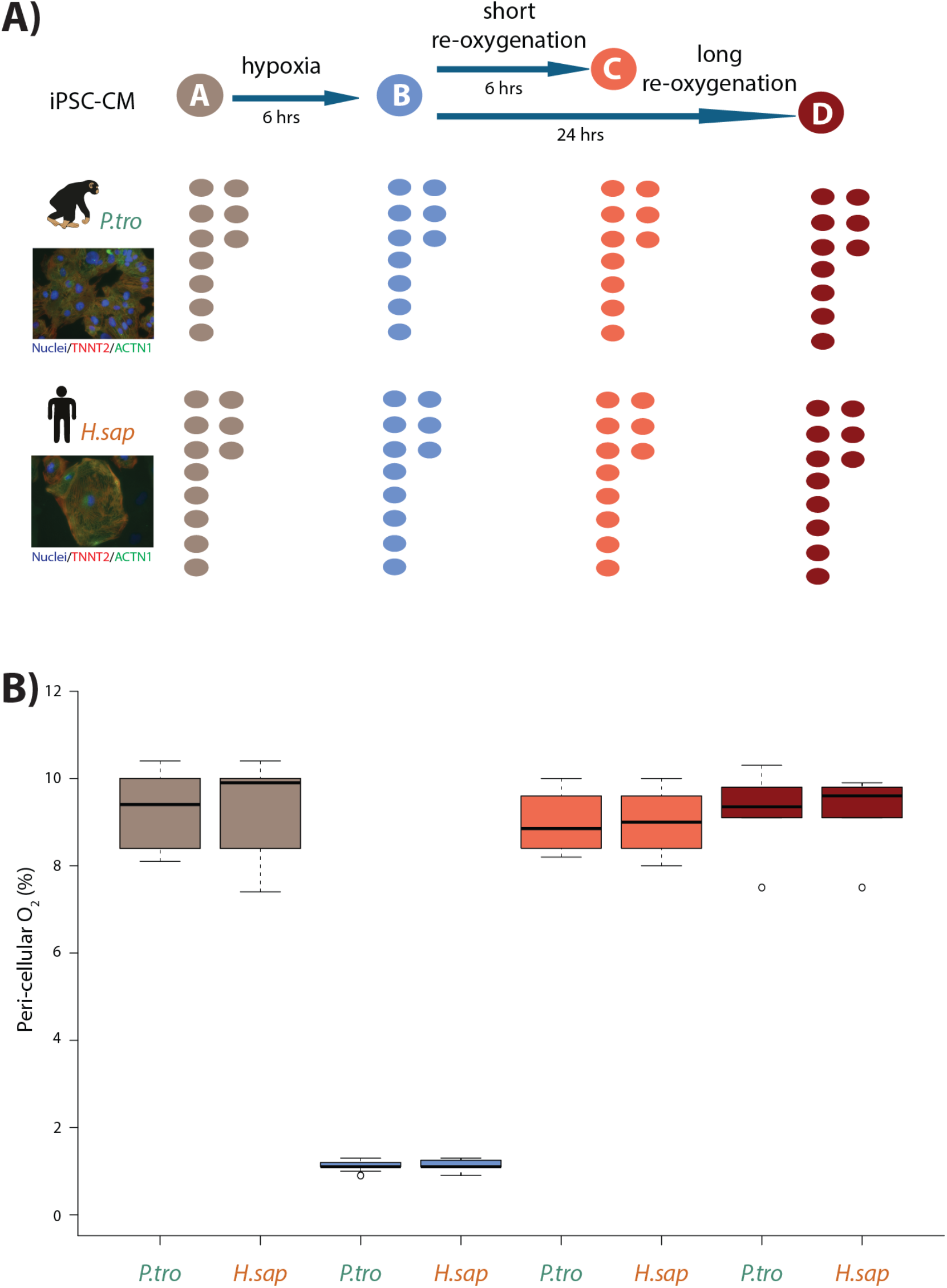
Induction of hypoxia in human and chimpanzee iPSC-CMs. **A)** Experimental design of the study. Cardiomyocytes were differentiated from iPSCs from eight human (*H.sap*), and seven chimpanzee (*P.tro*) individuals together with replicates from three individuals of each species. For the oxygen stress experiment iPSC-CMs in each species were cultured in normoxic conditions (10% O_2_ - condition A) for 6 hours prior to induction of hypoxia at 1% O_2_ for 6 hours (condition B). Following hypoxia, iPSC-CMs were re-oxygenated to 10% O_2_ for 6 hours (condition C), or 24 hours (condition D). **B)** Peri-cellular O_2_ levels measured at each stage of the experiment for each experimental batch.

After 30 days in culture, differentiated cells of both species express the cardiomyocyte markers cardiac troponin T (TNNT2), sarcomeric alpha-actinin (ACTN1), and Iroquois Homeobox 4 (IRX4) (Figure S1B). We determined the purity of differentiated cardiomyocytes in each culture independently, by measuring the proportion of cells expressing TNNT2 by flow cytometry (see Methods). Only samples with greater than 45% TNNT2-expressing cells were retained. Importantly, there is no difference in the median purity of the cardiomyocyte cultures between humans and chimpanzees (median purity in both species: 78%, Figure S3).

In order to mimic *in vivo* physiological oxygen levels experienced by cardiomyocytes in the heart, we cultured our cells at 10% oxygen, starting on the 25^th^ day of differentiation. Peri-cellular oxygen levels were measured non-invasively using an oxygen sensor applied to the inside sidewall of the cardiomyocyte cultures. Transferring the cardiomyocytes from atmospheric oxygen levels (21% O_2_) to 10% O_2_ did not seem to induce stress in the cultures (Figure S4A). In what follows, we consider 10% O_2_ to be the baseline normoxia condition (designated as condition A). In order to determine the response of the cultures to hypoxia and subsequent recovery to normoxia, we subjected the cardiomyocyte cultures to the following conditions (see Figure 1A for a schematic of the study design): We first lowered the O_2_ levels to 1% for 6 hours (condition B), we then re-oxygenated to 10% O_2_ for 6 hours of recovery in normoxic conditions (condition C), and 24 hours of recovery (condition D). Oxygen levels were monitored and recorded for each experimental batch (Figure 1B).

To confirm that perturbing the peri-cellular oxygen level affected cardiomyocytes from both species, we measured two phenotypes associated with oxidative damage. First, we determined whether damage to DNA is induced following hypoxic stress. Guanine is the nucleotide most prone to oxidation, and in general undergoes base excision repair. The repair products of the oxidative DNA lesions are subsequently released as 8-hydroxydeoxyguanosine (8-OHdG). The amount of 8-OHdG released therefore reflects both the amount of oxidative damage to DNA, and the efficiency of base excision repair. We observed an increase in the level of 8-OHdG following hypoxia and reoxygenation in both species (t-test; *P* = 0.002 in chimpanzees, and *P* = 0.01 in humans; Figure S5A). Within a condition, there is no difference in the amount of 8-OHdG released between species. Second, we determined the extent of lipid peroxidation by measuring the release of the isoprostane 8-iso-Prostaglandin F2α (8-iso-PGF2α), which is induced following ROS-mediated damage to cellular membrane phospholipids. We found an increase in 8-iso-PGF2α release in chimpanzees following hypoxia (*P* = 0.006), and a further increase following long-term reoxygenation (*P* = 0.03; Figure S5B). While there is no difference in 8-iso-PGF2α release between species within any condition, we do not find a significant increase in 8-iso-PGF2α release following hypoxia in humans (*P* = 0.2). This pattern may be explained by incomplete power to detect differences in 8-iso-PGF2α release either between time points or between species. Nevertheless, our observations are intriguing because 8-iso-PGF2α is known to be elevated in heart failure (Mallat et al. 1998; Wolfram et al. 2005), and is a risk marker for coronary heart disease (Schwedhelm et al. 2004).

### Characterizing the regulatory response to hypoxia in cardiomyocytes

We used RNA sequencing to characterize gene expression levels in all conditions, and study the regulatory response to hypoxia in the human and chimpanzee cardiomyocytes (see Methods). We processed the samples using a study design that was balanced with respect to a number of recorded potential technical confounders (Table S2). Following sequencing of the RNA, we mapped reads to primate orthologous exons (Figure S6), and filtered lowly-expressed genes to yield a final data set with expression measurements for 11,974 genes (see Methods). Within each condition, inter-species correlation in read counts is somewhat lower than intra-species variation, as expected (median Spearman’s correlation when comparing human samples = 0.97, when comparing chimpanzee samples = 0.98, and for human vs. chimpanzee samples = 0.92; Figure S7). Using the RNA-seq data we confirmed that genes known to be expressed in cardiomyocytes are expressed in both our human and chimpanzee samples, including genes involved in cardiac structure, ion channels, and adrenoreceptors (Figure S8).

As mentioned above, we included in our study differentiation replicates from a subset of samples (see Table S3 for details). We expect that gene expression data from pairs of replicates should be more similar to each other than to data from any other individual. We used this expected property of the data to account and correct the entire data set for unwanted technical variation (see Methods for more details). After accounting for unwanted variation, samples cluster by species and then by individual or condition (Figures S9). We note that one technical factor, the presence or absence of episomal reprogramming vectors (three human samples tested positive; Figure S10), remains a partial confounder with species. However, we confirmed that our conclusions are robust with respect to the inclusion of these three human samples (Figure S17 and Table S4).

To determine which genes respond to hypoxia, we analyzed the data from all four conditions using the framework of a linear model. The model included fixed effects for ‘species’, ‘condition’, a ‘species by condition interaction’, a random effect for ‘individual’, and four unwanted factors of variation as covariates (see Methods). For this analysis, we randomly selected one of each of the samples we had replicate data for. We were first interested in classifying genes into the following four categories within each species independently: genes that respond to hypoxia, genes that respond to short-term (6 hours) re-oxygenation following hypoxia, genes that respond to long-term (24 hours) re-oxygenation following hypoxia, and genes that differ between long-term reoxygenation and baseline normoxia. Of 11,974 expressed genes, we identified ~4,000 genes that respond to hypoxia at 10% FDR in each species, and a slightly higher number of genes whose expression has changed upon re-oxygenation (Figure 2; the results of all tests are in Table S4A).

**Figure 2:**
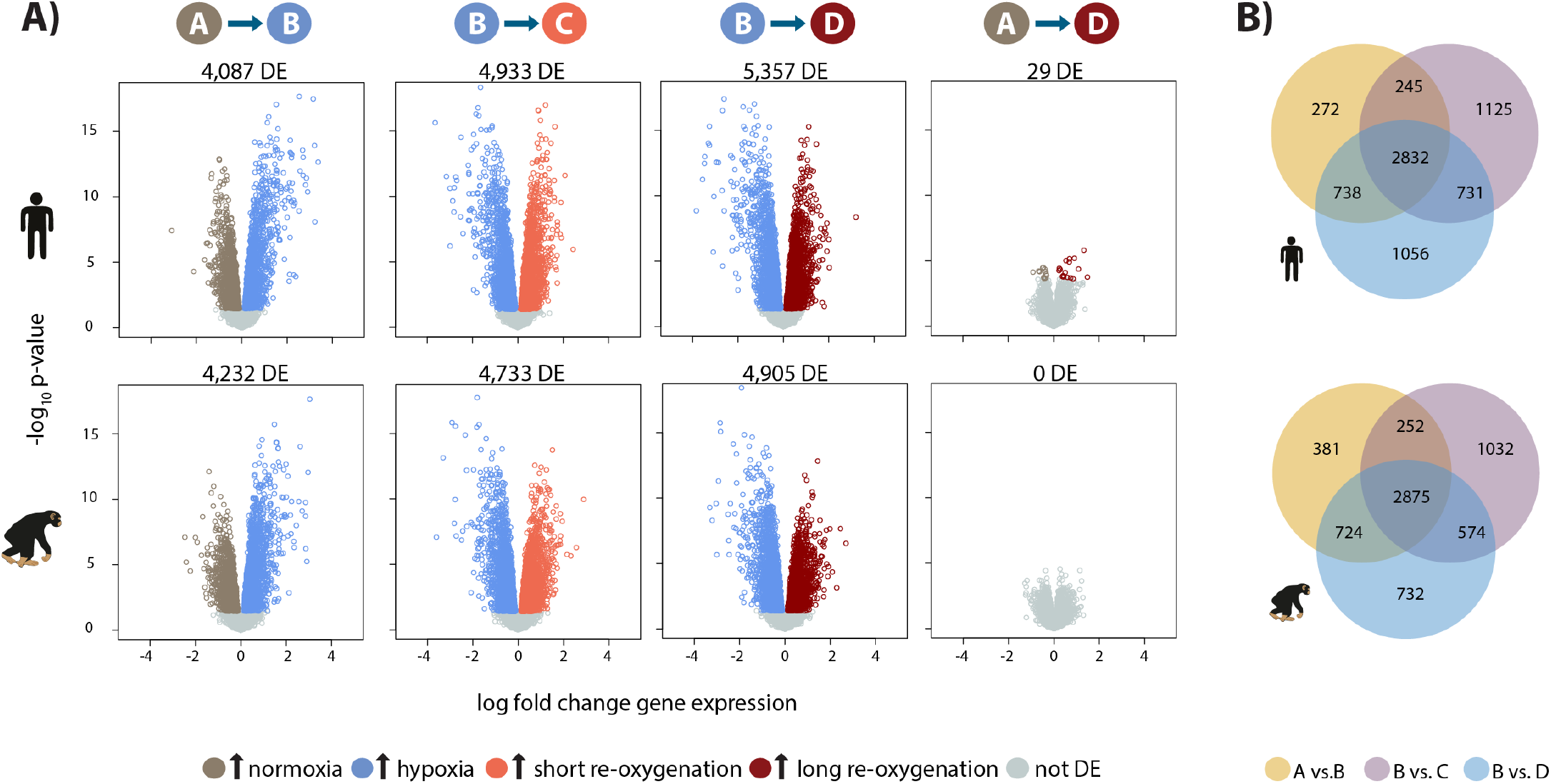
Hypoxia induces a gene expression response in humans and chimpanzees. A) Volcano plots representing genes that are differentially expressed (DE; 10% FDR) in pairwise comparisons across conditions in each species independently. In a comparison of A vs. B, genes that are up-regulated in hypoxia are represented in blue, and genes that are up-regulated in normoxia are represented in brown. Genes that are up-regulated in condition C are represented in coral, and genes that are up-regulated in D are represented in dark red. **B)** Overlap of genes that are differentially expressed in pairs of conditions in each species independently.

We then focused on inter-species gene expression differences within each condition, independently, and found that roughly half of all expressed genes are differentially regulated between species, regardless of the condition (at FDR of 10%; Table S4A, Figure S11A). Using this approach we were also able to identify hundreds of genes that are differentially expressed between species exclusively in a single condition, for example following hypoxia (Figure S11B). However, this approach does not provide strong evidence for true differences in the dynamic response to hypoxia between humans and chimpanzees, because of incomplete power to detect inter-species differentially expressed genes in any given condition. Thus, in order to determine the species-specificity of the global response to changing oxygen conditions we explicitly compared the effect size of expression change between pairs of conditions, for all genes, across species. Overall, there is a strong correlation in the global gene expression response to both hypoxia and re-oxygenation in humans and chimpanzees (median Spearman correlation = 0.78; sign test *P* < 10^−4^ for all comparisons; Figure 3), suggesting that the general response to changes in oxygen level is conserved in the two species. Genes that respond to either hypoxia or re-oxygenation in both species include *VEGFA* (a known hypoxia response gene), *TRPV1* (implicated in ischemia-reperfusion injury in the heart (Wang and Wang 2005)), and *DDX41* (implicated in stress survival regulation (Shih and Lee 2014)).

**Figure 3:**
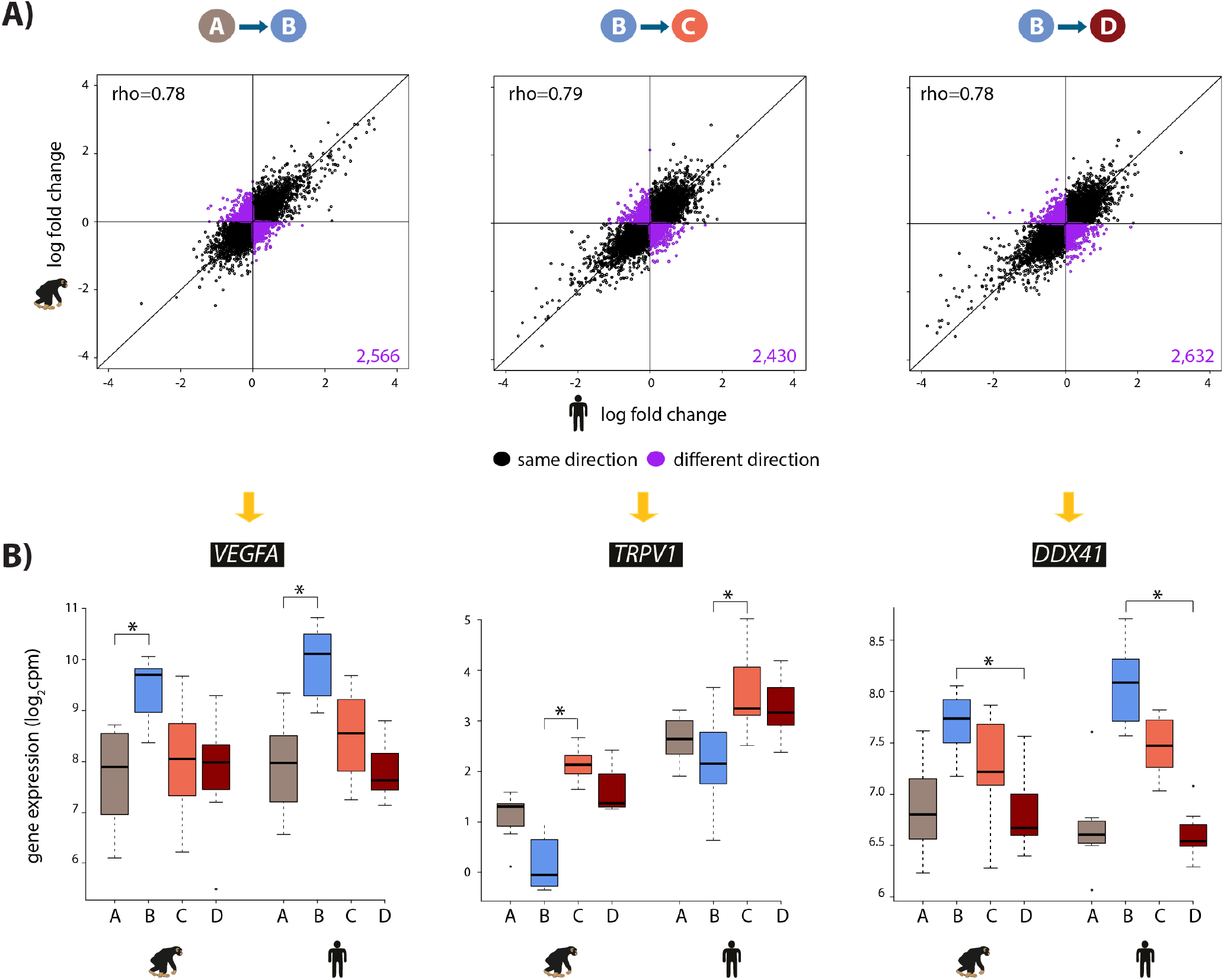
The hypoxic gene expression response is highly correlated across species. **A)** The log fold change in expression of 11,974 genes between pairs of conditions in humans on the x-axis, and chimpanzees on the y-axis. Genes whose expression changes in the same direction in both species are represented in black, and genes whose expression change direction differs across species are represented in purple. **B)** Examples of genes that are differentially expressed in both species in A vs. B (*VEGFA*), B vs. C (*TRpV1*), and B vs. D (*DDX41*) are shown. Asterisk denotes a statistically significant difference in expression between conditions (10% FDR).

### Species-specific transcriptional changes in response to hypoxia

The observation that the response to changes in oxygen level is generally conserved in the two species notwithstanding, we next focused on dynamic inter-species differences in our study. To do so, we used two approaches. First, we estimated a gene-specific interaction effect between species using the framework of the linear model described above. We identified 147 genes that responded to hypoxia in one species but showed little or no response in the other species, or that responded in both species but showed the opposite direction of effect (at FDR of 10%; Figure 4; Table 1). We did not find inter-species differences in the response to either the short or long reoxygenation treatments.

**Figure 4:**
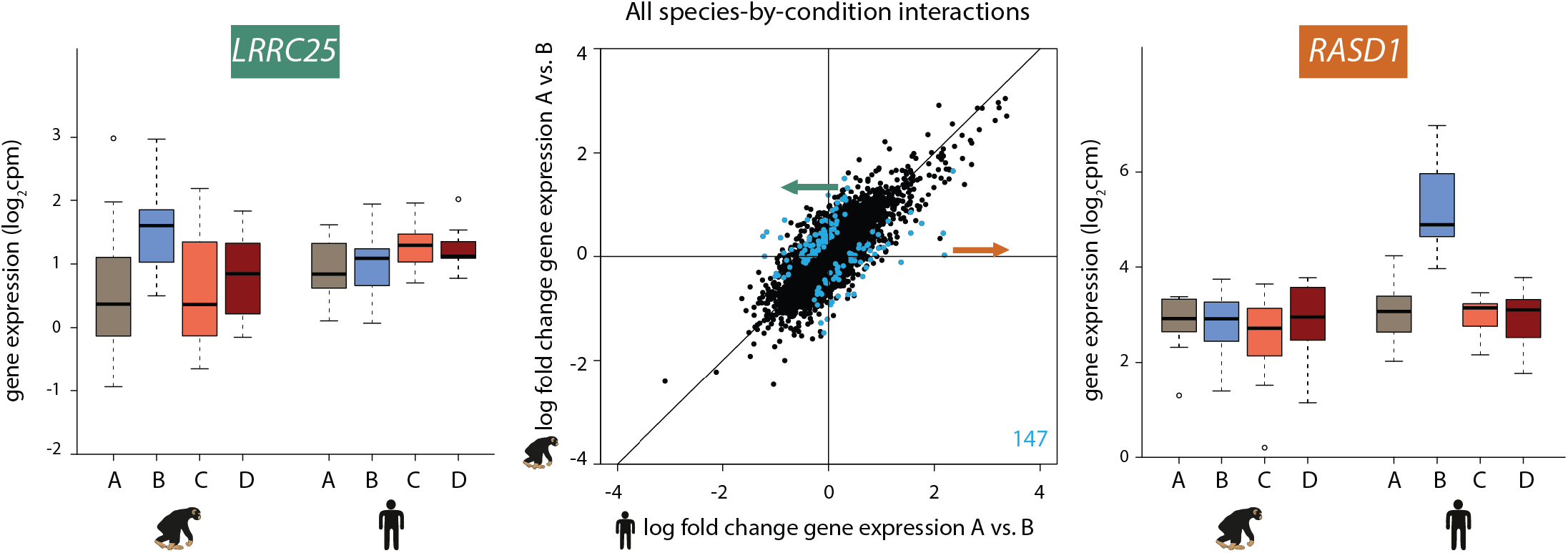
147 genes show a species-specific response following hypoxia. Middle panel: The log fold change in expression of 11,974 genes between normoxia (A) and hypoxia (B) in humans on the x-axis, and chimpanzees on the y-axis. Genes with a species-by-condition interaction are represented in blue. All species-by-condition interactions affected the hypoxic condition; therefore only a representative pairwise comparison is shown (A vs. B). An example of a gene that responds in chimpanzees only (*LRRC25*) is shown in the left panel, and an example of a gene that responds only in humans (*RASD1*) is shown in the right panel.

We did not find enrichment of particular pathways among the species-specific response genes, but this may not be surprising as stress response pathways are often regulated by a small number of key master regulator genes (Haynes et al. 2010; Li et al. 2011; Natarajan et al. 2013; Mahat et al. 2016; Quiros et al. 2017). However, several of the genes with significant species by condition interactions have functions related to G-protein signaling, TGF-β signaling, and metabolism (Figure S12). Our most significant interaction corresponds to the *RASD1* gene, which is up-regulated specifically in humans following hypoxia (Figure 4). This gene encodes a Ras GTPase, which activates G-protein signaling. Conversely, the *LRRC25* gene responds to hypoxia specifically in chimpanzees, and has been found to inhibit NF-κβ signaling (Feng et al. 2017) (Figure 4).

Directly modeling interaction effects with small numbers of samples is an underpowered approach. In order to side-step the challenge of incomplete power when performing multiple pairwise comparisons, we used a second approach; a joint Bayesian model, to classify genes based on their expression levels during the course of the hypoxia-re-oxygenation experiment (see Methods). Four gene clusters were empirically determined to explain the predominant expression patterns in the data (lowest BIC and AIC after testing 1-15 clusters; Figure S13). Using this approach we categorized 9,414 genes as not responding to hypoxia in either humans or chimpanzees (nonresponse genes), 1,920 genes that respond to hypoxia in both species (conserved response genes), 430 genes that respond to hypoxia in chimpanzees only (chimpanzee-specific response genes), and 199 genes that respond to hypoxia only in humans (human-specific response genes; Figure 5 and Table 2). It is notable that there is no prevalent pattern of genes responding specifically to re-oxygenation in either species, which suggests that the expression of most genes returns to baseline by the end of the experiment. We do not identify additional gene expression patterns even when we increase the number of clusters.

**Figure 5:**
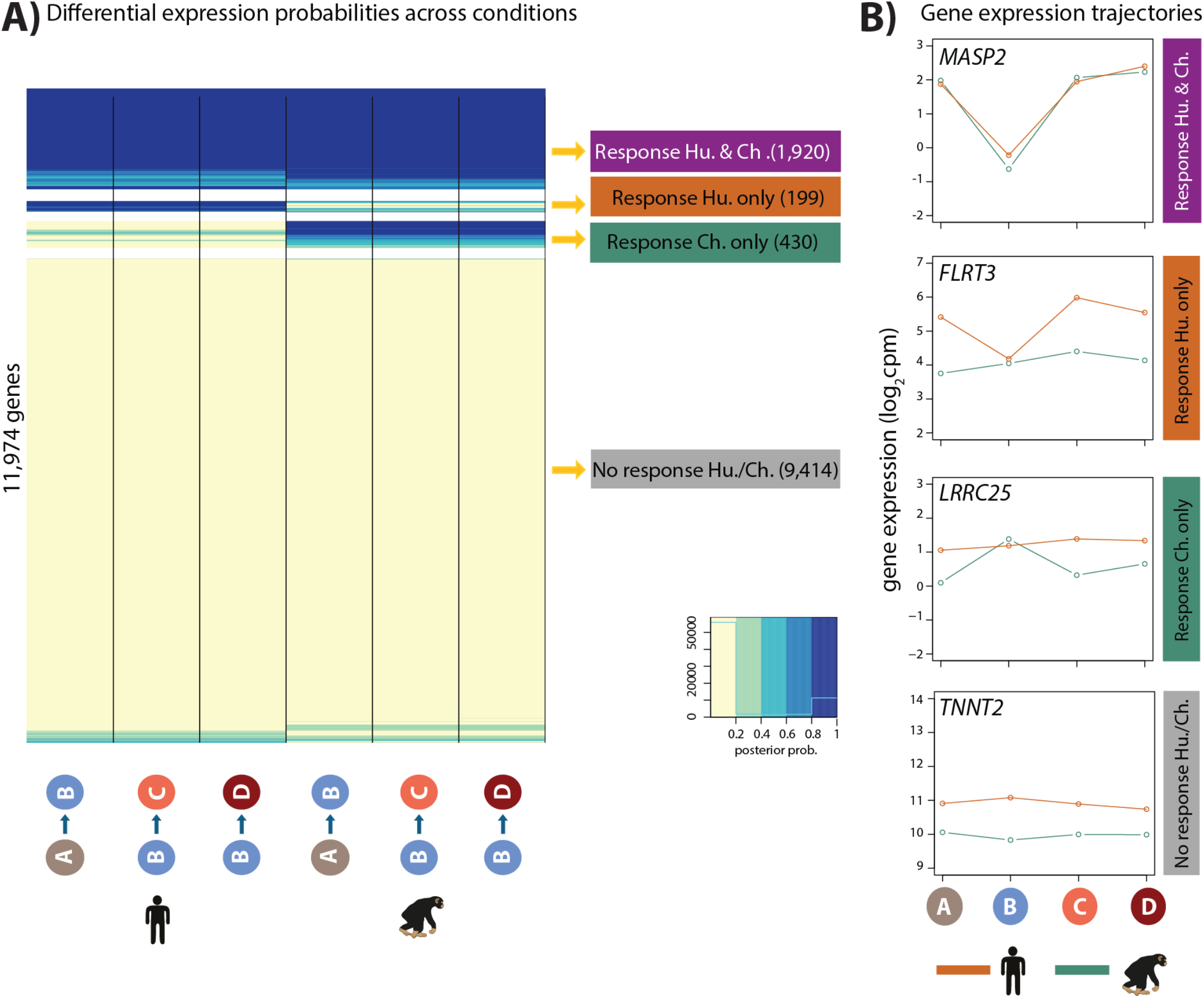
Hypoxia induces four gene expression trajectories across species. **A)** Posterior probabilities of genes being differentially expressed across pairs of conditions. Genes are categorized based on their posterior probabilities: genes with a p < 0.5 across contrasts are designated as “Non-response genes” (grey), genes with a p > 0.5 across contrasts are designated as “Conserved response genes” (magenta), genes with p > 0.5 in human comparisons only are designated as “Human-specific response genes” (orange), and genes with p > 0.5 in chimpanzee comparisons only are designated as “Chimpanzee-specific response genes” (green). **B)** Examples of genes belonging to each of the four categories. Mean expression levels per species are plotted.

### Properties of conserved hypoxic response genes

To confirm that our approach identifies meaningful response genes, we considered the overlap of genes assigned to our four response categories with a set of genes previously identified to respond to hypoxia in human, and a more evolutionary distant primate, the rhesus macaque (Zhao et al. 2018). As expected, genes previously found to respond to hypoxia are enriched among genes assigned to the conserved response category in our study, and depleted among genes assigned to the ‘non-response’ category (Chi-squared test; *P* < 10^−15^ in both; Figure S14A).

We next wanted to determine what distinguishes these four gene expression response clusters. To do so, we first tested whether the genes in the response categories are enriched for particular pathways, using a background set of all expressed genes. In the non-response gene category, there is a significant enrichment in KEGG pathways related to the heart (e.g. dilated cardiomyopathy, hypertrophic cardiomyopathy, arrhythymogenic right ventricular cardiomyopathy and adrenergic signaling in cardiomyocytes, 10% FDR; Figure S14B). In the conserved response category, various signaling pathways related to sensing the external environment, and responding to oxygen are significantly enriched including HIF1α, MAPK and FOXO1. There are no significantly enriched pathways in the species-specific gene response categories. Given the apparent enrichment of cardiovascular genes in the non-response category, we explicitly tested the contribution of a set of cardiovascular-associated genes to the response to hypoxia (see Methods). Indeed, we found that there is a depletion of genes implicated in cardiovascular development and disease amongst the genes that respond to hypoxia in both species (Chi-squared test; *P* = 8.3×10^−6^; Figure 6A).

**Figure 6:**
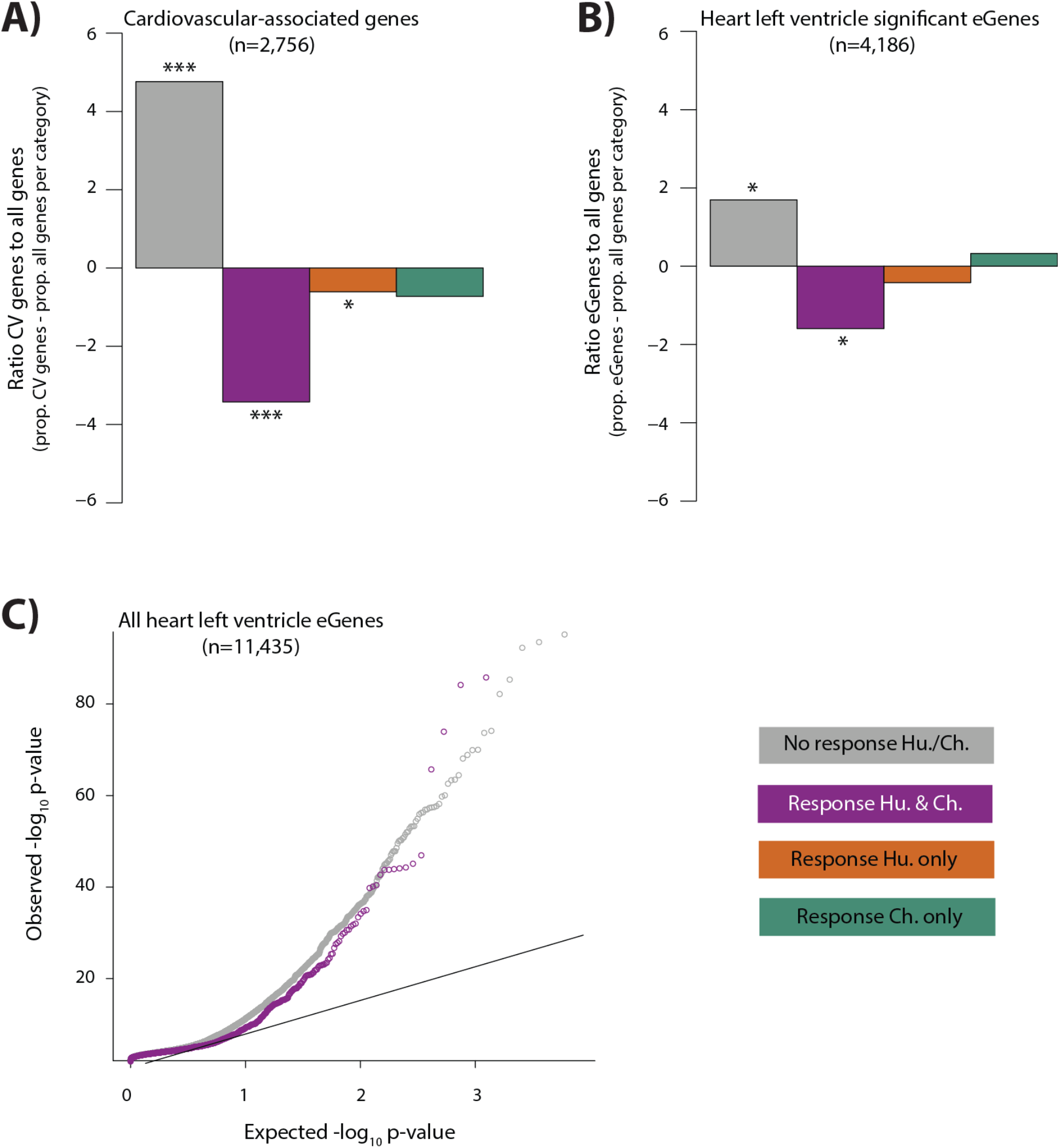
Cardiovascular-associated genes and eGenes in heart tissue are depleted in conserved hypoxic response genes. **A)** The proportion of cardiovascular-associated (CV) genes (Cardiovascular GO Annotation Initiative) in each response category relative to the proportion of all genes within a response category: non-response (grey), conserved response (magenta), human-specific response (orange), and chimpanzee-specific response (green). Asterisk denotes a significant difference between the proportion of cardiovascular genes in each response category, and the proportion of all genes within a response category (Chi-squared test; \**P* < 0.05, \*\*\**P* < 0.0005). **B)** The proportion of heart left ventricle eGenes (GTEx Consortium) in each response category, relative to the proportion of all genes in each category. **C)** QQ-plot representing all eGenes identified in heart left ventricle that overlap the non-response category (grey), and conserved response category (magenta).

It has been suggested that genetic variants that associate with gene expression levels (eQTLs), may mediate disease phenotypes (Emilsson et al. 2008; Albert and Kruglyak 2015; Yao et al. 2015; Gamazon et al. 2018). In order to test the contribution of eQTLs to the response to stress, a phenotype that is likely to provide insight into disease, we overlapped our four response gene categories with genes whose expression level is associated with genetic variants in human heart tissues (eGenes; see Methods). We observe a depletion of eGenes in the conserved response category, when compared to all expressed genes within each category, using data from the heart left ventricle (Chi-squared test; *P* = 0.01), heart right atrial appendage (*P* = 1.4×10^−5^), and iPSC-derived cardiomyocytes (*P* = 6.4×10^−5^; Figure 6B and Figure S15). The depletion corresponds to a difference in the contribution of eGenes to the non-response and conserved response categories (*P* < 10^−15^; Figure 6C). The pattern of depletion of eGenes among conserved response genes is also observed across 12 other tested tissues; however the magnitudes of the effect differ between tissues (Figure S15B). We found that human-specific response genes are also depleted of eGenes when we considered all eGenes identified in at least 1 of the 14 tested tissues (*P* = 7.1×10^−5^).

As we observed depletion of eQTLs found in healthy individuals among the conserved response genes in our study, we next considered eQTLs found among CVD patients. To do so, we investigated the contribution of eQTLs identified in left ventricle heart tissue from patients undergoing aortic valve replacement surgery pre- and post-cardioplegic arrest and ischemia, to our hypoxia response categories (Stone et al. 2018). Again, we observed a depletion of eGenes in the conserved response category for both pre- and post-ischemia eGenes (*P* = 0.03 and *P* =0.006 respectively; Figure S16A). Interestingly, we observed the opposite pattern when we considered genes that are differentially expressed between pre- and post-ischemia samples (*P* = 0.14 for nonresponse genes and *P* = 0.17 for conserved response genes; Figure S16B).

## Discussion

Studying the human response to stress in an evolutionary context can potentially provide insight into disease susceptibility, incidence, aetiology, and response to treatment. In order to understand human adaptation and susceptibility to oxygen deprivation in the heart, we developed a cell-culture model to study the response to, and recovery from, hypoxia in iPSC-CMs from humans and chimpanzees. Using this system we were able to control exposure to changing oxygen levels in both species and study the ensuing *in vitro* response.

We found that, in humans and chimpanzees, the expression of ~4,000 genes (about a third of all expressed genes) is altered following six hours of hypoxic stress. Many of these genes return to baseline expression levels within 24 hours of re-oxygenation. The response to hypoxia is highly conserved in the two species with 1,920 genes responding similarly in humans and chimpanzees (75% of all genes that respond in at least one species). There have not been many comparative studies of functional perturbations in primates to provide us with broad context, but the conservation in response to hypoxia in our study is much greater than the conservation in immune response to infection between humans and chimpanzees (Barreiro et al. 2010). This suggests that the response to oxygen is a fundamentally conserved process across species, unlike the rapidly-evolving response to pathogens.

Conserved hypoxic response genes correspond to signaling pathways related to oxidative stress and hypoxia including the FOXO1 and HIF1 signaling pathways. However, genes responding to hypoxia are significantly depleted for known cardiovascular-associated genes. This suggests that hypoxic stress response genes are expressed, and active in multiple tissues. This is supported by the fact that we observe that conserved response genes are also depleted for eGenes identified in tissues unrelated to the heart. These results suggest that there is less tolerance for genetic variability that results in variation in the expression of genes that are necessary for eliciting a response to stress. Cellular stress, including oxidative stress, can contribute to cellular damage and lead to disease pathology (Giordano 2005; Sack et al. 2017).

A common notion is that genetic variants that modulate gene expression levels (eQTLs) are important in mediating disease phenotypes (Emilsson et al. 2008; Albert and Kruglyak 2015; Yao et al. 2015; Gamazon et al. 2018). Thousands of genetic variants that associate with various phenotypes, including disease presentation, have been identified through genome-wide association studies (GWAS). Given that most of these variants are located within non-coding regions of the genome, it is thought that integrating GWAS data with eQTL data will help to identify genes that are relevant to the phenotype of interest (Hormozdiari et al. 2016; Zhu et al. 2016). The observation that up to half of GWAS-identified variants are also eQTLs in at least one tissue (Consortium et al. 2017), provided some measure of support for this notion. However, there are several lines of evidence, which indicate that the relationship between eQTLs and complex disease may be relevant mainly to diseases that manifest late in life, and are therefore unlikely to have an effect on fitness: (i) Highly constrained genes with missense and protein-truncating variants, that might be expected to contribute to disease, are depleted for eQTLs but enriched for GWAS variants (Lek et al. 2016). (ii) Most eQTLs are shared across a large number of tissues and are expected to have broad functional effects, which are therefore unlikely to be highly deleterious (Consortium et al. 2017). (iii) eQTLs identified in other primates are often also eQTLs in human (Tung et al. 2015; Jasinska et al. 2017); suggesting that, across species, eGenes can tolerate the accumulation of associated mutations, which perturb their regulation. Together, these findings suggest that most eQTLs may be neutral. In this study, we performed perturbation experiments to determine the consequences on gene expression across species. Our results, which show a depletion of eQTLs in genes that respond to stress across species, support the notion that eQTLs alone may not give immediate insight into disease phenotypes.

Genetic variants that modulate gene expression levels only in response to direct perturbation (response QTLs) are more likely to be informative in disease. Indeed, the association between GWAS variants is more pronounced in response QTLs than naïve QTLs (Barreiro et al. 2012; Alasoo et al. 2018). There are currently a limited number of data sets that allow for a systematic investigation of this association. However, using an available data set of heart tissue from CVD patients, we again observed a depletion of eQTLs in stress response genes. Our results suggest that eQTLs correspond to a set of genes that are largely distinct from genes, which respond to stress, or which are relevant to disease. Indeed, genes that are differentially expressed between pre- and post-ischemia samples show the opposite pattern to that of eQTLs in our data i.e. they are enriched in genes that respond to stress across species. While many eQTLs act in *cis*, it has been suggested that *trans*-eQTLs are more likely to associate with complex traits (Westra et al. 2013; Consortium et al. 2017). However, we are currently underpowered to confidently identify these variants and determine their relevance to stress and disease.

In addition to quantifying gene expression levels in response to hypoxia, we measured oxidative stress phenotypes in our comparative cardiomyocyte system. The baseline level of DNA oxidation damage is similar in humans and chimpanzees, and increases following recovery from hypoxia in both species. The baseline level of lipid peroxidation is also similar in humans and chimpanzees, and shows a trend towards increased levels during the course of the experiment in both species; however this increase is only significant in chimpanzees. These findings are in line with a study of oxidative stress markers in blood from ten male humans and ten male chimpanzees, which showed that there is no significant difference in the levels of 8-OHdG between species, but there is significantly elevated 8-iso-PGF2α levels in chimpanzees compared to humans (Videan et al. 2009). Another comparative study on the effects of hypoxia in human and rhesus macaque cardiomyocytes demonstrates that the secreted metabolome is highly correlated between species after 24 hours of hypoxia (Zhao et al. 2018), suggesting an additional layer of regulatory conservation.

### Inter-species differences in response to oxygen perturbation

Although the overall correlation in the response to hypoxia is high between humans and chimpanzees, a stringent interaction analysis identified 147 genes with species-specific expression in the hypoxic condition.

The most significant species-specific response gene is *RASD1*, which is similarly expressed in humans and chimpanzees in normoxic conditions but is significantly up-regulated after hypoxia only in humans. *RASD1* was found to be up-regulated in samples from patients with ischemic disease compared to patients with dilated cardiomyopathy (heart damage despite normal blood flow), and non-failing hearts (Liu et al. 2015). These results suggest that aberrant *RASD1* expression could be specifically involved in the response to oxygen deprivation, and the pathogenesis of ischemic heart disease. In addition, the *RAI1-PEMT-RASD1* region is a replicated, genome-wide significant locus for coronary artery disease (CAD) (McPherson and Tybjaerg-Hansen 2016). It is unclear what pathway, related to the CAD phenotype, is affected by the *RASD1* locus (Khera and Kathiresan 2017). RASD1 is an activator of G-protein signaling (Cismowski et al. 2000), and is thought to contribute to the stress response (Sato and Ishikawa 2010). Indeed, G-proteins are important sensors of the environment at the cell membrane and mediate a signaling cascade to initiate an intra-cellular response to external stimuli. The potential role of RASD1 in mediating the response to hypoxia is unclear as RASD1 inhibits the secretion of the cardiac hormone ANF following volume overload experiments in the heart (McGrath et al. 2012), yet hypoxia induces ANF release from hearts (Baertschi et al. 1986). *RASD1* is one of several species-specific response genes related to G-protein signaling; other examples include *ARRDC2, RASL11B, ARL6 RASSF1, GTPBP4, RAB3A, RHOF, SYDE2*, and *CNKSR1*.

In fact, there are several genes that respond in a species-specific manner, which belong to similar pathways, or perform similar functions. For example, multiple genes (*ACVR2A, SNIP1, JUNB, SMAD4, TGFBR2, SMAD6, FGF9*) are related to TGF-β signaling. TGF-β is induced following myocardial infarction, and mediates the development of fibrosis (Bujak and Frangogiannis 2007). It is tempting to speculate that differences in the expression of these genes under oxygen stress, could contribute to the fibrotic heart phenotype observed in chimpanzees. Several of these genes have been implicated in heart physiology and disease suggesting that they could be relevant to the phenotype (Galvin et al. 2000; Wang et al. 2005; Alfonso-Jaume et al. 2006; Tseng et al. 2009; Itoh et al. 2016; Lu et al. 2016; Dogra et al. 2017).

Many species-specific response genes are involved in post-transcriptional layers of the gene regulatory cascade including RNA modifications (*METTL14*), RNA folding (*DDX20*), splicing (*CCDC49*), nuclear-cytoplasmic transport (*NUP214*), and protein degradation (*CBLL1, UBQLN4, TRIM13, DNAJB2)*. This suggests that additional inter-species differences may emerge in processes downstream of transcription. For example, METTL14 deposits the N^6^-methyladenosine RNA modification, which has been implicated in the stabilization of mRNA molecules following hypoxia (Fry et al. 2017). Intriguingly, differences in N^6^-methyladenosine levels have been reported between primates (Ma et al. 2017). Alternative splicing is another mechanism for inter-species differences between humans and chimpanzees, and, interestingly, one gene that undergoes differential splicing between species is the *GSTO2* gene, which is protective against oxidative stress (Calarco et al. 2007).

Genes that respond to oxygen deprivation in a species-specific manner could have phenotypic consequences as rapid changes in gene expression in response to stress can lead to evolutionary adaptation (Lopez-Maury et al. 2008; de Nadal et al. 2011). This mechanism has been implicated in mediating inter-species differences in epithelial cancer incidence as coordinated gene expression differences between humans and chimpanzees have been observed in fibroblasts subjected to serum starvation (Pizzollo et al. 2018). Conversely, it has been suggested that species-specific gene responses to stress across divergent yeast species may be adaptive, or, more likely, compensated by the response of related genes, or reflective of biological noise (Tirosh et al. 2011). The latter could explain the overall inter-species similarity in the cellular and transcriptional response, and the fact that genes peripherally related to similar pathways show species-specific differences.

In summary, to date there have been few well-powered studies investigating the evolution of the stress response in primates. Here we measured the genome-wide transcriptional response to a universal cellular stress, oxygen deprivation, across species in a CVD-relevant cell type. We find that the cellular and transcriptional response is largely similar across species; however there are hundreds of genes that respond in a species-specific manner.

## Methods

### Samples

We used eight biological replicates (individuals) from human, and seven from chimpanzee. In addition, technical replicates (independent cardiomyocyte differentiation and oxygen stress experiments) from three human and three chimpanzee individuals were used to estimate unwanted factors of variation in the data. All iPSC lines were derived from fibroblasts. 13 iPSC lines have been described and characterized previously (Gallego Romero et al. 2015; Burrows et al. 2016; Pavlovic et al. 2018; Ward et al. 2018). Two additional iPSC lines are first described and characterized in this study (H22422 and H25237). All sample metadata is included in Table S1.

### Differentiating cardiomyocytes from iPSCs

Feeder-independent iPSCs were maintained at 70% confluence on Matrigel hESC-qualified Matrix (354277, Corning, Bedford, MA, USA) at a 1:100 dilution. Cells were cultured in Essential 8 Medium (A1517001, ThermoFisher Scientific, Waltham, MA, USA) with Penicillin/Streptomycin (30002Cl, Corning) at 37°C, 5% CO_2_ and atmospheric O_2_. Cells were passaged every 3-4 days with dissociation reagent (0.5 mM EDTA, 300 mM NaCl in PBS), and seeded with ROCK inhibitor Y-27632 (ab12019, Abcam, Cambridge, MA, USA).

iPSC-CM differentiations were largely performed based on the protocol described by Burridge *et al*. (Burridge et al. 2014). Importantly, the same differentiation protocol was used in both species. iPSCs cultured for 10-50 passages were seeded in 4 × 10 cm Matrigel-coated culture dishes until 70-100% confluent (Days -4/-3). The optimum cell density for efficient differentiation depended on the individual iPSC line. On Day 0, 6 μM of the GSK3 inhibitor, CHIR99021 trihydrochloride (4953, Tocris Bioscience, Bristol, UK) was added to the cultures in 12 ml Cardiomyocyte Differentiation Media [500 mL RPMI1640 (15-040-CM ThermoFisher Scientific), 10 mL B-27 Minus Insulin (A1895601, ThermoFisher Scientific), 5 mL Glutamax (35050-061, ThermoFisher Scientific), and 5 mL Penicillin/Streptomycin)], and a 1:100 dilution of Matrigel. 24 hours later, on Day 1, fresh Cardiomyocyte Differentiation Media, supplemented with 6 μM CHIR99021 was added to the cultures. Chimpanzee iPSCs, in general, were more sensitive to the addition of CHIR99021 hence the reduction from the optimal 12 μM CHIR99021 for 24 hours as described in Burridge *et al*., to 6 μM for 48 hours. The GSK3 inhibitor was removed with the addition of fresh Cardiomyocyte

Differentiation Media 24 hours later on Day 2. After 24 hours, on Day 3, 2 μM of the Wnt signaling inhibitor Wnt-C59 (5148, Tocris Bioscience), diluted in Cardiomyocyte Differentiation Media, was added to the cultures. The media was replaced with 2 μM Wnt-C59 in Cardiomyocyte Differentiation media 24 hours later, on Day 4. Cardiomyocyte Differentiation Media was replaced on Days 5, 7, 10 and 12. Spontaneously beating cells appear on Days 7-10.

To remove non-cardiomyocytes from the cultures, iPSC-CMs were purified by metabolic selection. 10 mL of glucose-free, lactate-containing media (Purification Media) [500 mL RPMI without glucose (11879, ThermoFisher Scientific), 106.5 mg L-Ascorbic acid 2-phosphate sesquimagenesium salt (sc228390, Santa Cruz Biotechnology, Santa Cruz, CA, USA), 3.33 ml 75 mg/ml Human Recombinant Albumin (A0237, Sigma-Aldrich, St Louis, MO, USA), 2.5 mL 1 M lactate in 1 M HEPES (L(+)Lactic acid sodium (L7022, Sigma-Aldrich)), and 5 ml Penicillin/Streptomycin] was added on Day 14. Purification Media was replaced on Days 16 and 18.

On Day 20, iPSC-CMs were dissociated with 4 mL 0.05% Trypsin-EDTA solution (25-053-Cl, ThermoFisher Scientific) for ~10 min, and quenched with double the volume of Cardiomyocyte Maintenance Media [500 mL DMEM without glucose (A14430-01, ThermoFisher Scientific), 50 mL FBS (S1200-500, Genemate), 990 mg Galactose (G5388, Sigma-Aldrich), 5 mL 100 mM sodium pyruvate (11360-070, ThermoFisher Scientific), 2.5 mL 1 M HEPES (SH3023701, ThermoFisher Scientific), 5 mL Glutamax (35050-061, ThermoFisher Scientific), 5 mL Penicillin/Streptomycin]. This change in carbohydrate source shifts metabolism away from glycolysis towards aerobic mitochondrial respiration, which is the predominant pathway used to generate energy in adult cardiomyocytes *in vivo*. A single cell suspension was generated by straining the cells through a 100 μm nylon mesh cell strainer three times, and once through a 40 μm mesh strainer. 1.5 million iPSC-CMs were plated per well of a Matrigel-coated 6-well plate in 3 mL Cardiomyocyte Maintenance Media. iPSC-CMs for each of the four conditions were plated on separate 6-well plates.

### Culturing iPSC-CMs

iPSC-CMs were matured in culture for a further 10 days. Cardiomyocyte Maintenance Media was replaced on Days 23, 25, 27, 28 and 30.

While cells are typically cultured *in vitro* at atmospheric oxygen levels (21% O_2_), this oxygen level is not experienced by mammalian cells *in vivo* - arterial oxygen levels are ~13%, and levels drop to 5-10% within tissues (Brahimi-Horn and Pouyssegur 2007; Carreau et al. 2011; Jagannathan et al. 2016). We therefore chose to culture our cells at 10% O_2_, which falls within this physiological range. On Day 25, iPSC-CMs cultured at atmospheric oxygen levels, were transferred to an oxygen-controlled incubator (HERAcell 150i CO_2_ incubator, ThermoFisher Scientific) representing physiological oxygen levels (10% O_2_). Oxygen levels are maintained through displacement of oxygen by nitrogen.

To allow further maturation, and to synchronize iPSC-CM beating, iPSC-CMs were pulsed with the IonOptix C-Dish & C-Pace EP Culture Pacer from Day 27 until the end of the oxygen perturbation experiment. Cells were pulsed at a voltage of 6.6 V/cm, frequency of 1 Hz, and pulse frequency of 2 ms.

### Immunocytochemistry

iPSC-CM samples were dual-stained in two independent staining reactions.

#### Stain 1

Cells were fixed with 3-4% paraformaldehyde for 15 min at room temperature and then washed three times with PBS. Cells were permeabilized with 0.3% Triton X-100 for 10 min prior to another three washes with PBS. Cells were blocked with 5% BSA for 30 min and then incubated with primary antibodies diluted in 5% BSA O/N at 4°C. Anti-Cardiac Troponin T rabbit polyclonal antibody (ab45932, Abcam) was added at a 1:400 dilution. Anti-alpha-Actinin(Sarcomeric) mouse monoclonal antibody (A7811, Sigma) was added at a 1:500 dilution. After primary antibody incubation, cells were washed with 0.1% Tween-20 in PBS three times. Secondary antibodies, donkey anti-Rabbit IgG Alexa Fluor 594 (A21207, ThermoFisher Scientific), and Donkey antiMouse IgG Alexa Fluor 488 (A21202, ThermoFisher Scientific), were added at a 1:1000 dilution in 2.5% BSA for 1 hr at room temperature. Cells were washed three times with PBS prior to counter-staining with Hoechst for 10 min.

#### Stain 2

Cells were fixed with 3-4% paraformaldehyde for 15 min at room temperature and washed three times with PBS. Cells were not permeabilized or blocked. Primary antibodies were diluted in Permeabilization Buffer (FOXP3/Transcription Factor Staining Buffer Set, 00-5523, ThermoFisher Scientific), and incubated O/N at 4°C. Anti-IRX4 rabbit polyclonal antibody (ab123542, Abcam) at a 1:200 dilution, and Cardiac Troponin T Monoclonal Antibody (MA5-12960 clone 13-11, ThermoFisher Scientific) at a 1:200 dilution, were used. Cells were washed three times with Permeabilization Buffer after primary antibody incubation. Secondary antibodies were added at a 1:1000 dilution in Permeabilization Buffer for 1 hr at room temperature. Donkey anti-Mouse IgG Alexa 594 (A21203, ThermoFisher Scientific), and Donkey anti-Rabbit IgG Fluor 488 (A21206, ThermoFisher Scientific), were used. Cells were washed three times with PBS prior to counter-staining with Hoechst for 10 min.

### Flow cytometry

iPSC-CMs were dissociated with 0.05% Trypsin-EDTA solution and quenched with four times the volume of Cardiomyocyte Maintenance Media. In order to obtain a single cell suspension, cells were strained twice through a 100 μm strainer, and once through a 40 μm strainer. 1 million cells were stained with Zombie Violet Fixable Viability Kit (423113, BioLegend) for 30 min at 4°C prior to fixation and permeabilization (FOXP3/Transcription Factor Staining Buffer Set, 00-5523, ThermoFisher Scientific) for 30 min at 4°C. Cells were stained with 5 μl PE Mouse Anti-Cardiac Troponin T antibody (564767, clone 13-11, BD Biosciences, San Jose, CA, USA) for 45 min at 4°C. Cells were washed three times in permeabilization buffer and re-suspended in autoMACS Running Buffer (130-091-221, Miltenyi Biotec, Bergisch Gladbach, Germany). Several negative controls were used in each flow cytometry experiment: 1) iPSCs, which should not express TNNT2, 2) an iPSC-CM sample that has not been labeled with viability stain or TNNT2 antibody, and 3) an iPSC-CM sample that is only labeled with the viability stain.

10,000 cells were captured and profiled on the BD LSRFortessa Cell Analyzer. Several gating steps were performed to determine the proportion of TNNT2 positive cells: 1) Cellular debris was removed by gating out cells with low granularity on FSC versus SSC density plots, 2) From this population, live cells were identified as the violet laser-excitable, Pacific Blue dye-negative population, 3) Two populations of TNNT2 positive cells were identified within the set of live cells: one conservative gate selected high-intensity TNNT2 cells, and a lenient gate included TNNT2 cells with a range of intensities. Both gates were created so as to exclude any cells that overlap the profiles of the negative control samples. iPSC-CM purity is reported as the proportion of TNNT2-positive live cells.

### Oxygen stress experiment

Several pilot experiments were performed to determine the optimal oxygen conditions to initiate a gene expression response in iPSC-CMs. The expression of stress response genes is induced within 6 hours of hypoxia (Figure S4B), and cell damage (lactate dehydrogenase activity) is evident after 24 hours (Figure S4C). It is noteworthy that culturing the iPSC-CMs in media with a glucose carbohydrate source, instead of galactose, does not elicit a stress response following hypoxia (Figure S4). Given that we were interested in determining the early transcriptional response to hypoxia, prior to the induction of cell death, we chose the 6-hour time-point in our subsequent experiments.

Oxygen stress experiments were conducted on Day 31 or 32 after the initiation of differentiation. At the start of the experiment (total elapsed time = 0 hrs), one plate (A) remained at 10% O_2_, while three plates (B, C and D) were transferred to an oxygen-controlled cell culture incubator set at 1% O_2_. After six hours, plates A and B were harvested, while plates C and D were transferred back to 10% O_2_ (elapsed time = 6 hrs). Plate C was harvested six hours after the end of the hypoxic incubation (elapsed time = 12 hrs). Plate D was harvested 24 hours after the end of the hypoxic incubation (elapsed time = 30 hrs). Ten batches of oxygen stress experiments were performed. For each of the four conditions, iPSC-CMs were harvested by manual scraping, flash-frozen as cell pellets, and stored at −80°C, together with the cell culture media from each sample, until further processing.

Oxygen levels in the cell culture media of a representative iPSC-CM sample were measured during the course of the oxygen perturbation experiment, in each experimental batch. An oxygen sensitive sensor was applied to the inner wall of a well of a 6-well plate (SP-PSt3-NAU-D5-YOP, PreSens Precision Sensing GmbH, Regensburg, Germany). Oxygen levels were measured non-invasively through the wall of the cell culture plate using a Polymer Optical Fiber (NWDV29, Coy, Grass Lake, MI, USA), and a Fiber Optic Oxygen Meter (Fibox 3 Transmitter NWDV16, Coy).

### Lactate Dehydrogenase Activity Assay

Lactate dehydrogenase (LDH) activity levels were measured by colorimetric determination of NAD reduction to NADH using the Lactate Dehydrogenase Activity Assay Kit (MAK066, Sigma-Aldrich). 5 μl of cell culture media was assayed as per the manufacturer’s instructions. Each sample was assayed in triplicate. LDH activity is reported as the difference in NADH levels measured at the start of the enzymatic reaction, and 25 min after the addition of the substrate. Enzyme activity is calculated relative to a linear standard curve.

### Oxidative DNA damage assay

8-OHdG levels were measured by competitive enzyme-linked immunoassay using the OxiSelect Oxidative DNA Damage ELISA Kit (STA-320, Cell Biolabs Inc.). Levels were measured in 50 μl of cell culture media, in duplicate, according to the manufacturer’s instructions. Samples were processed on three species-balanced 96-well plates. 8-OHdG was quantified relative to a standard curve using 4- and 5-parameter logistic models implemented in the drc package in R. Final 8-OHdG release is reported as four measurements: A (A-A), B (B-A), C (C-B) and D (D-B). For the three individuals with replicate experiments in each species, mean values from both experiments are reported.

### Lipid peroxidation assay

Excreted 8-iso-PGF2α was measured by competitive enzyme-linked immunoassay using the OxiSelect 8-iso-Prostaglandin F2α ELISA kit (STA-337, Cell Biolabs Inc., San Diego, CA, USA). Levels were measured in 55 μl of cell culture media, in duplicate, according to the manufacturer’s instructions. Samples were processed on three species-balanced 96-well plates. 8-iso-PGF2α was quantified relative to a standard curve using 4- and 5-parameter logistic models implemented in the drc package in R. Final 8-iso-PGF2α release is reported as four measurements either relative to the baseline, condition A, or the hypoxic condition i.e. A (A-A), B (B-A), C (C-B) and D (D-B). For the three individuals with replicate experiments in each species, mean values from both experiments are reported.

### RNA-seq library preparation and sequencing

RNA was extracted from ~1.5 million cells from 84 iPSC-CM samples representing 15 individuals. Extractions were performed in ten species-balanced batches using the ZR-Duet DNA/RNA extraction kit (D7001, Zymo, Irvine, CA, USA). All four conditions from one human and one chimpanzee individual were extracted per batch (one batch had three individuals). RNA concentration and quality was measured using the Agilent 2100 Bioanalyzer. RIN scores were greater than 7.5 for all samples (human median: 9.1, chimpanzee median: 9.2).

RNA-seq libraries were prepared from 250 ng of RNA in three species-balanced batches using the Illumina TruSeq RNA Sample Preparation Kit v2 (RS-122-2001 & -2002, Illumina). Libraries in each batch were multiplexed together to generate four pools for sequencing. Each pool was sequenced 50 base pairs, single-end on the HiSeq4000 according to the manufacturer’s instructions. Pools 1,3,4 were sequenced on three lanes (24 samples per pool), and Pool 2 was sequenced on two lanes (12 samples in the pool).

### RNA-seq data processing

RNA-seq data quality was determined by FastQC (http://www.bioinformatics.babraham.ac.uk/projects/fastqc/). Sequencing adapters were trimmed, and sequencing reads from each species aligned to their respective genome (hg19 or panTro3) using TopHat2 (version 2.0.11)(Kim et al. 2013). The number of mapped sequencing reads is similar across species and conditions (median human A: 53,547,009, median human B: 40,872,328; median human C: 40,635,553, median human D: 42,041,844; median chimpanzee A: 42,258,525; median chimpanzee B: 33,054,882; median chimpanzee C: 28,792,150, median chimpanzee D: 36,903,485)

We quantified gene expression levels at orthologous meta-exons from 30,030 Ensembl genes from hg19, panTro3 and rheMac3 (Blekhman et al. 2010) using featureCounts within subread (version 1.4.6) (Liao et al. 2014). Genes were filtered to only include those on the autosomes. Log_2_-transformed counts per million were calculated using edgeR (Robinson et al. 2010). Lowly-expressed genes were filtered such that only genes with a mean log_2_ cpm > 0 across samples were retained.

Prior to differential expression analysis, unwanted factors of variation were estimated in the RNA-seq data using RUVSeq (Risso et al. 2014). As we have two replicate samples from six of the individuals (three in each species), the RUVs function for estimating the factors of unwanted variation using replicate samples was used. This approach takes advantage of the fact that replicate samples have constant covariates of interest. We tested different numbers of unwanted factors of variation (k values) until the data clustered best by our biological factors of interest i.e. species, individual and condition. Four factors of unwanted variation were thus selected.

To assess data quality we performed Principal Component Analysis (PCA) on the RUVs-normalised log_2_ cpm expression values. We correlated known biological and technical factors with the first six PCs.

### Differential expression analysis

The TMM-voom-limma pipeline was used to identify differentially expressed genes between species and conditions. Filtered read counts from a randomly selected replicate were taken forward in this analysis. Normalization factors were used to scale the raw library size to the effective library size of each sample using the trimmed mean of M-values (TMM) implemented in edgeR (Robinson et al. 2010). The mean-variance relationship was removed using precision weights in voom (Law et al. 2014). Correlation between samples from the same individual was estimated using the duplicateCorrelation function.

A linear model was fitted for expression values of each gene using limma (Smyth 2004). To compare pairwise differential expression, the species, condition and a species-by-condition interaction were modeled as fixed effects, and individual as a random effect. The four unwanted factors of variation determined by RUVs are modeled as covariates. In order to obtain more precise gene-wise variability estimates, we used empirical Bayes moderation, which takes information across all genes into account. We used contrast tests in limma to identify genes that are differentially expressed between conditions within each species, genes that are differentially expressed between species at each condition, and species-by-condition interactions for conditions B, C, and D. We corrected for multiple testing at each gene using the Benjamini & Hochberg false discovery rate (FDR) (Benjamini and Hochberg 1995). Genes with FDR-adjusted *P* values of < 0.1 are considered to be differentially expressed.

### Identification of gene expression trajectories

In order to cluster genes by their gene expression trajectories during the course of the experiment, all data was jointly modeled using a Bayesian Hierarchical model across pairwise differential tests implemented in Cormotif (Wei et al. 2015). To identify gene expression trajectories (correlation motifs), expression levels were compared across conditions within each species (1: human A vs. B, 2: human B vs. C, 3: human B vs. D, 4: chimpanzee A vs. B, 5: chimpanzee B vs. C, 6: chimpanzee B vs. D). TMM-normalised cpm values were used as input data. The Bayesian information criterion (BIC) and Akaike information criterion (AIC) was used to select the best model when varying the number of correlation motifs from 1 to 15 at a set seed. The BIC and AIC were minimized when four correlation motifs were modeled. The posterior probability of differential expression for each gene, in each pairwise test, is plotted in the heatmap. Gene sets corresponding to the four correlation motifs were defined as having posterior probabilities > or < 0.5 (motif 1 (non-response): p < 0.5 in contrasts 1,2,3,4,5,6; motif 2 (chimpanzee-specific response): p < 0.5 in contrasts 1,2,3 and p > 0.5 in contrasts 4,5,6; motif 3 (conserved response): posterior probability > 0.5 in contrasts 1,2,3,4,5,6; motif 4 (human-specific response): p > 0.5 in contrasts 1,2,3 & p < 0.5 in contrasts 4,5,6).

### Comparison of our data to that collected in human and rhesus macaque iPSC-CMs

The set of 187 genes that were previously found to respond to hypoxia in both human and rhesus macaque iPSC-derived cardiomyocytes were overlaid with our four Cormotif gene expression response categories (Zhao et al. 2018). 164 of these genes are expressed in our data. Results are presented as a difference in the proportions between genes that respond to hypoxia in humans and rhesus macaques overlapping our four response categories, and all genes in each of our four response categories. Statistical significance between proportions is assessed using a Chi-squared test.

### Gene ontology analysis

The gene sets belonging to each of the four response categories were investigated for common pathway enrichment using the KEGG database (Kanehisa et al. 2017), within the DAVID genomic annotation tool (Huang da et al. 2009b; Huang da et al. 2009a). Enrichment was calculated relative to the set of all 11,974 expressed genes. Multiple testing was performed by the Benjamini-Hochberg method. Pathways enriched at 10% FDR are considered to be significant.

### Cardiovascular gene analysis

A set of 5,010 genes implicated in cardiovascular development or disease (BHF-UCL gene association file) was obtained from the Cardiovascular Gene Ontology Annotation Initiative (https://www.ebi.ac.uk/GOA/CVI). The 2,756 genes that are expressed in our study were overlapped with our four response categories. Results are presented as a difference in the proportions between cardiovascular genes overlapping our four response categories, and all genes in each of our four response categories. Statistical significance between proportions is assessed using a Chi-squared test.

### eQTL analysis

For the overlap of our response categories with eGenes from healthy individuals, the list of eGenes in 14 GTEx tissues was downloaded from v7 in the GTEx portal (www.gtexportal.org). eGenes were selected at 5% FDR in each tissue. eGenes from iPSC-derived cardiomyocytes were obtained from Banovich *et al*. (10% FDR) (Banovich et al. 2018).

For the overlap of our response categories with eQTLs identified pre- and post-ischemia, an RNA-seq study of 114 patients undergoing aortic valve replacement surgery was used (Stone et al. 2018). In this study, samples of heart left ventricle were obtained for each individual pre- and post- cardioplegic arrest/ischemia. Lists corresponding to all eGenes (5% FDR) identified in males and females combined within the pre-ischemic state (496 expressed in our data), all eGenes in the post-ischemic state (416 expressed), and differentially expressed genes (5% FDR) between pre- and post-ischemia (6,571 in 46 females, and 6,572 in 68 males) were used.

Results are presented as the difference in proportions between all eGenes overlapping our response categories, and all genes in each response category; or as a difference in the proportions of differentially expressed genes overlapping our response categories, and all genes in each response category. Statistical significance between proportions is assessed using a Chi-squared test.

## Supplemental material

### S1 Supplemental material

Document containing supplemental figures 1-17 and supplemental tables 1-4.

### Table 1

Table with species-by-condition interaction genes.

### Table 2

Table with all genes assigned to the four gene expression response categories.

### Data Access

All RNA-seq data have been deposited in the Gene Expression Omnibus (www.ncbi.nlm.nih.gov/geo/) under accession number GSE117192.

## Acknowledgements

We thank all members of the Gilad lab for helpful discussions, Kristen Patterson for experimental assistance, and the Genomics Core Facility at the University of Chicago for sequencing the RNA-seq libraries. We thank The Genotype-Tissue Expression (GTEx) Project, supported by the Common Fund of the Office of the Director of the National Institutes of Health, and by NCI, NHGRI, NHLBI, NIDA, NIMH, and NINDS, for providing data. The data used for the analyses described in this manuscript were obtained from the GTEx portal v7 on May 24^th^ 2018. M.C.W is supported by an EMBO Long-Term Fellowship (ALTF 751-2014), and the European Commission Marie Curie Actions. This work was funded by NIH grants GM120167 and HL139447.

## Contributions

M.C.W and Y.G conceived and designed the study. M.C.W performed experiments and analysed data. M.C.W and Y.G wrote the manuscript. Y.G oversaw the study.

